# Distinct taxonomic groups of sponge symbionts have the biosynthetic capacity for the production of ether lipids

**DOI:** 10.64898/2026.06.23.733905

**Authors:** Catarina Loureiro, Michelle A. Schorn, Diana X. Sahonero Canavesi, Asimenia Gavriliidou, Vasilis Gerovasileiou, John van der Oost, Laura Villanueva, Marnix H. Medema, Detmer Sipkema

## Abstract

The marine sponge holobiont, composed of the sponge host and its microbial symbionts, is a known source of abundant and diverse ether lipids (ELs). Apart from their structural role in the cytoplasmic membrane of archaea and some bacteria, ELs have often been linked to signaling functions and defense against pathogens. Despite the relevance of ELs, their biosynthesis, as well as the identity of their producers, remain elusive. Here, we report the analysis of potential ether lipid producing genes and gene clusters, detected in marine sponge metagenomes as well as public sponge genomes. We show that the sponge holobiont has the capacity to synthesize ELs via several pathways, and suggest the ability of the sponge holobiont to synthesize ELs under different O2 levels. Finally, targeted lipidome analysis confirmed that ELs are present in the lipid profiles of all of the studied sponge holobiont samples, and indicates that the biosynthesis of the plasmalogens detected is likely restricted to the sponge host itself, based on the detected hydrocarbon chain lengths. This work provides a basis for the challenging quest to decipher intricate EL biosynthesis in marine sponges and their associated microbes.

## Introduction

Marine sponges, comprising the phylum Porifera, are primitive multicellular organisms that inhabit widely varied aquatic habitats as heterotrophic sessile filter feeders (Van Soest *et al*., 2012; McMurray *et al*., 2016; Webster and Thomas, 2016). High microbial abundance sponges host highly diverse communities of microbial symbionts in their tissues that are often dominated by Proteobacteria, Actinobacteria, Acidobacteriota, Chloroflexota, Cyanobacteria and the sponge-specific candidate phylum Poribacteria (Hentschel *et al*., 2003; Thomas *et al*., 2016; Podell *et al*., 2019). These microbial symbionts have been proposed to play a role in promoting host health (Fan *et al*., 2012; Webster and Thomas, 2016; Engelberts *et al*., 2020). In parallel, the sponge host provides shelter and nutrition to its microbial community (Rix *et al*., 2020). Marine sponges are considered extremely rich sources of natural products (NPs) with notable bioactivities, and their symbiotic associations are linked to an extended metabolic repertoire, with many sponge-holobiont-derived NPs demonstrated to be of symbiotic bacterial origin (Agarwal *et al*., 2017; Calcabrini *et al*., 2017; Morita and Schmidt, 2018; Carroll *et al*., 2019; Schorn *et al*., 2019). The enzymes responsible for the biosynthesis of bacterial NPs are, in most cases, encoded by biosynthetic gene clusters (BGCs) (Medema *et al*., 2015). BGCs originating from the marine environment, and more specifically those encoded by bacterial sponge symbionts, often display unique features (Loureiro *et al*., 2022).

Many sponges harbor highly abundant bacterial membrane lipids with fatty acids linked through ether bonds to a backbone containing phosphate or mono/disaccharide (glyco) polar head groups, instead of the more common ester-linked phospholipids (Dembitsky *et al*., 1989; Gramzow *et al*., 1989; Djerassi and Lam, 1991; Quévrain *et al*., 2012) **(Figure 1)**. These ELs can be of two types, namely alkyl-acyl glycerophospholipid with an alkyl ether bond, and alkenyl-acyl glycerophospholipid with an alkenyl ether bond, also known as 1-alkyl-1⍰-enyl, vinyl ether or plasmalogen. Plasmalogens have been suggested to have functions related to protection against pathogens, signaling, colonization, maintenance of the membrane structure, acid tolerance and protection against reactive oxygen species (Braverman and Moser, 2012; Vítová *et al*., 2021; Goldfine, 2022; Mu *et al*., 2024). Sponge, and also coral, holobiont ELs are often linked to pathogen defense through their antimicrobial activity (Müller *et al*., 2004; Quinn *et al*., 2016; Mohsenian Kouchaksaraee *et al*., 2020; Rubio-Portillo *et al*., 2020), and they have also been reported to show anticancer activity (Alam et al. 2001; Dyshlovoy et al. 2022; Ben-Califa et al. 2012; Fedorov et al. 2009). Thus, sponge ELs are considered to be interesting sponge-holobiont-derived NPs with potential biotechnological applications. The exact producers of sponge-holobiont associated ELs, and their biosynthetic pathways, are still unknown.

**Figure 1:**
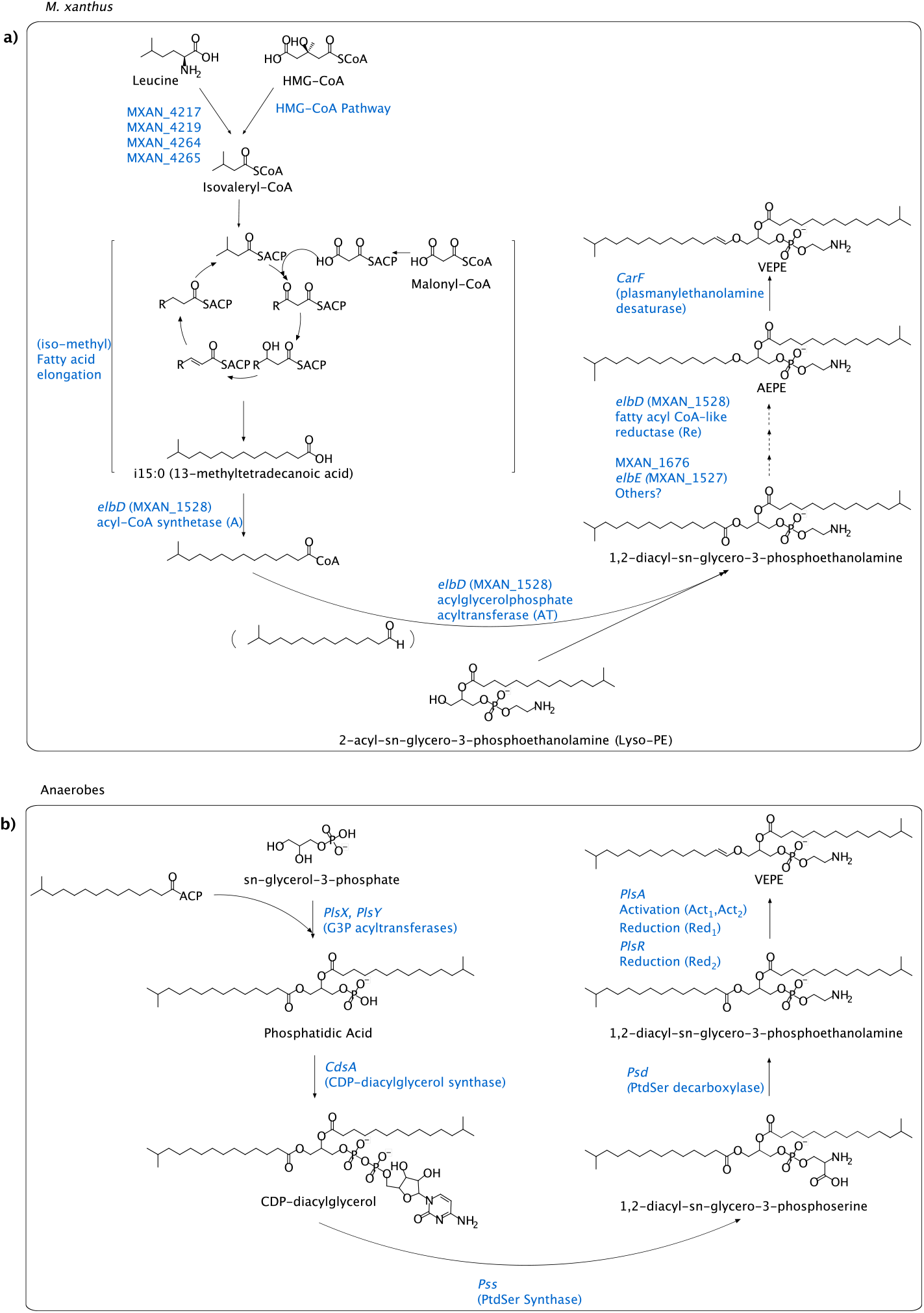
Proposed bacterial biosynthesis of plasmalogens. a) Proposed AEPE/VEPE biosynthesis in *M. xanthus*; b) Proposed VEPE biosynthesis pathway in anaerobes. Adapted from (21,27,28).

Outside of the sponge holobiont, bacterially produced ELs have been detected in cultures of anaerobic bacteria (i.e, sulfate-reducing bacteria and members of the order Clostridiales) (Goldfine, 2017), and also in those of specific aerobic bacteria (i.e., Acidobacteria and Myxobacteria) (Řezanka *et al*., 2012; Sinninghe Damsté *et al*., 2014), and their biosynthesis has been recently elucidated in selected taxa. Aerobic synthesis of (alkyl and alkenyl) ELs has been observed in the myxobacterium *Myxococcus xanthus* and involves the *elbB-elbE* BGC, as well as a desaturase-encoding gene *carF* (Lorenzen *et al*., 2014; Gallego-García *et al*., 2019). An alternative *M. xanthus* pathway involving an alkylglycerone-phosphate synthase (AGPS) produces minor amounts of plasmalogen EL (Lorenzen *et al*., 2014).

*ElbD*-homologs, as well as *agps* gene homologs, have been previously reported in Acidobacteriota Subdivision 4 (SD 4), but have yet to be conclusively linked to the biosynthetic pathway of acidobacteriotal ELs (Sinninghe Damsté *et al*., 2018; Sahonero-Canavesi *et al*., 2022). Acidobacteriota, specifically those belonging to SD 4 isolated from soil environments (Kielak *et al*., 2016), have been shown to synthesize alkyl ether bonds in their membrane lipids, which contain *iso*-C_15:0_ as an abundant fatty acid and features the phosphocholine headgroup in the phospholipid fraction (Sinninghe Damsté *et al*., 2014). An elbD homologue of poribacterial origin has also previously been reported by Lorenzen et al. (Lorenzen et al. 2014), although there is no record of ether lipid biosynthesis in this phylum to this date.

In addition, oxygen-sensitive EL biosynthetic pathways have been demonstrated in strict and facultative anaerobic microbes. For the synthesis of plasmalogens, a multi-subunit complex encoded by the *pls* operon, composed of the PlsA and PlsR enzymes, directly converts a diacyl precursor ester to a plasmalogen vinyl ether through reduction and dehydration *(Jackson et al., 2021)*. In addition, a homolog of the *plsA/R* genes, a glycerol ester reductase (Ger) has recently been shown to be involved in the synthesis of alkyl ether lipids in anaerobic bacteria (Sahonero-Canavesi *et al*., 2022).

Despite the fact that two alternative EL biosynthetic pathways have been observed, little is known about the biological advantages of producing ELs and the identity of the microbial EL producers, specifically in sponges where they are quantitatively important.

For this study, we set out to investigate the capacity of the sponge holobiont to produce ELs and which biosynthetic pathways are utilized in a dataset of high microbial abundance (HMA) sponges from varying depths and locations. We leverage state-of-the-art computational tools developed for detection and comparison of biosynthetic gene clusters, as well as key biosynthetic genes of the aerobic and anaerobic EL pathways. We establish a detailed overview of the diversity and distribution of *elb*-like BGCs, and link these BGCs, as well as standalone genes, to their putative bacterial hosts. We combine these findings with paired lipid analysis to confirm the presence of mono alkyl and alkenyl (plasmalogens) glycerophospholipids in all analyzed sponges. Finally, we identify host sponge gene homologs that could be responsible for the production of long chain plasmalogens.

## Materials and Methods

### Sponge collection

This study included samples from *Aplysina aerophoba (Chaib De Mares et al., 2018), Geodia barretti* collected in Canada (CAN) and Norway (NOR), *Petrosia ficiformis* (Loureiro *et al*., 2022), as well as *Aciculites cribophora, Smenospongia* sp., *Caminus* sp. and *Aplysina* sp. samples (Peters *et al*., 2023), all collected and processed in previous studies (Table S1).

### Total Lipid extraction and lipid analysis

For lipid analysis, tissue material from *A. aerophoba* (all samples), *G. barretti* (CAN & NOR) (gb1, gb2, gb5, gb8, gb126), and *P. ficiformis* (Pf5, Pf6, Pf9, Pf11) was freeze-dried and acid hydrolyzed as described previously (Sahonero-Canavesi *et al*., 2022). The extracts were processed with diazomethane and silylated with pyridine and with N,O-bis(trimethylsilyl)trifluoroacetamide (BSTFA) to derivatize alcohol groups. Monoalkyl glycerol ethers (MGE) and dimethylacetals (DMA) were identified by retention time and their GC-MS fragmentation (mass-to-charge, m/z, 205 base peak for alkyl ethers, m/z 75 base peak for dimethylacetals and their corresponding molecular ion).

### Total DNA extraction, metagenomic sequencing and sequence processing

#### Sponge eDNA extraction

*A. aerophoba (Chaib De Mares et al., 2018), G. barretti* (CAN & NOR) and *P. ficiformis* sponge samples, as well as *A. cribophora, Smenospongia* sp., *Caminus* sp. and *Aplysina* sp. samples (Peters *et al*., 2023) were extracted as in previous studies for generation of eDNA for short read Illumina sequencing.

*A. cribophora, Smenospongia* sp., *Caminus* sp. and *Aplysina* sp. samples were also subjected to two rounds of multiplexed Nanopore sequencing. For this, DNA was extracted using the MagAttract® HMW DNA kit (Qiagen), following the Disruption/Lysis of Tissue protocol, followed by the Manual Purification of High-Molecular Weight Genomic DNA from Fresh or Frozen Tissue protocol. Sponge pieces were weighed after thawing and squeezed to remove RNALater but were not rinsed.

#### Metagenomic sequencing, assembly and binning

*A. aerophoba (Chaib De Mares et al., 2018), G. barretti* (CAN & NOR) and *P. ficiformis* sponge samples, as well as *A. cribophora, Smenospongia* sp., *Caminus* sp. and *Aplysina* sp. samples (Peters *et al*., 2023) were sequenced as in previous studies, using the Illumina HiSeq PE150 and Illumina NovaSeq platforms.

For the first round of Nanopore sequencing of *A. cribophora, Smenospongia* sp., *Caminus* sp. and *Aplysina* sp., the Ligation Sequencing Kit SQK-LSK109 (Oxford Nanopore Technologies), the NEBNext Companion Module for Oxford Nanopore Technologies Ligation Sequencing kit (New England Biolabs®), and NBD104 barcodes (Oxford Nanopore Technologies) were used to create the sequencing libraries, following the Ligation sequencing gDNA - native barcoding (SQK-LSK109 with EXP-NBD104) protocol. The second sequencing library was made using the same kits, except new barcodes from the NBD114 kit (Oxford Nanopore Technologies) were used and the Ligation sequencing gDNA - native barcoding (SQK-LSK109 with EXP-NBD104 and EXP-NBD114) protocol was followed. Samples of 200 ng of each of five sponge libraries were combined for the final sequencing library. The first round of sequencing was performed on an already used, flushed flow cell (R9.4.1), with approximately 590 pores available. The second round of sequencing was done on a new flow cell (R9.4.1), with approximately 1301 pores available.

Quality control, trimming and adapter removal of *A. aerophoba, G. barretti* (CAN & NOR), *P. ficiformis* Illumina reads have been described in (Loureiro *et al*., 2022) and for *A. cribophora, Smenospongia* sp., *Caminus* sp. and *Aplysina* sp. in (Peters *et al*., 2023). Metagenomic read assembly of *A. aerophoba, G. barretti, P. ficiformis* has been described in our previous study (Loureiro *et al*., 2022). Hybrid assemblies of *A. cribophora, Smenospongia* sp., *Caminus* sp. and *Aplysina* sp. Illumina and Nanopore reads were performed here using metaSPAdes v3.12 with – only-assembler option.

Metagenome binning and MAG classification of *A. aerophoba, G. barretti* and *P. ficiformis* was done in (Loureiro *et al*., 2022). For *A. cribophora, Smenospongia* sp., *Caminus* sp. and *Aplysina* sp. reads were binned and classified following the same strategy used for *A. aerophoba* contigs (Loureiro *et al*., 2022). MAG abundance in host samples was predicted using the MetaWRAP quant module (Uritskiy *et al*., 2018). The obtained MAGs were de-replicated using dRep (Olm *et al*., 2017) v2.5.4 with default parameters for primary clustering and secondary clustering using parameters --S_algorithm gANI --S_ani 0.95 (Olm *et al*., 2020).

#### EL-like biosynthetic pathway detection and annotation

*A. aerophoba, G. barretti* (CAN & NOR), *P. ficiformis* BGCs were predicted using antiSMASH v5 *(Blin et al., 2019)* in our previous study (Loureiro *et al*., 2022), and *A. cribophora, Smenospongia* sp., *Caminus* sp. and *Aplysina* sp. BGC prediction followed the same process here. EL-like BGCs were identified using CORASON (Navarro-Muñoz *et al*., 2020) v1, default parameters, with gb8_2 contig 859 gene 9 as a query and MIBiG BGC0000871.1 as the reference BGC. BiG-SCAPE (Navarro-Muñoz *et al*., 2020) v1.0.1 was run on all EL-like BGCs using parameters --mix -v -- mode auto --mibig --cutoffs 0.5 --include_singletons to generate a preliminary set of Gene Cluster Families (GCFs). This set was used for curation and trimming of EL-like BGCs, i.e. exclusion of putatively truncated BGCs, and trimming based on manually curated putative borders (Table S2) using Python scripts available at https://github.com/CatarinaCarolina/sponge_ether_lipid. A final set of GCFs and representative BGCs was constructed using BiG-MAP (Pascal Andreu *et al*., 2021) with parameters -tg 0.1 -c 0.3 for the family module. AMP-domain amino acid sequences of each representative BGC (Fig. 2), as well as all BGCs (Fig. S4), were used to construct two multiple sequence alignments (MSA) with Muscle (Edgar, 2004) v3.8.31. FastTree (Price *et al*., 2010) v2.1.11 was used to process these MSAs into phylogenetic trees, which were visualised with iTOL (Letunic and Bork, 2019). Prediction of functional class and substrate specificity prediction based on adenylation domain active-site-specificity-conferring residues was conducted with AdenylPred v1 (Robinson *et al*., 2020) and PARAS v1 (https://github.com/BTheDragonMaster/paras) with default settings. PARAS-residues v0.0.3 was used to extract and align protein sequences of A-domain active site and Stachelhaus residues. MAG features, i.e., presence of BGCs in MAGs, MAG taxonomy, MAG abundance in sample, were integrated by processing GTDB-Tk and BiG-MAP outputs with in-house Python scripts available at https://github.com/CatarinaCarolina/sponge_ether_lipid.

**Figure 2:**
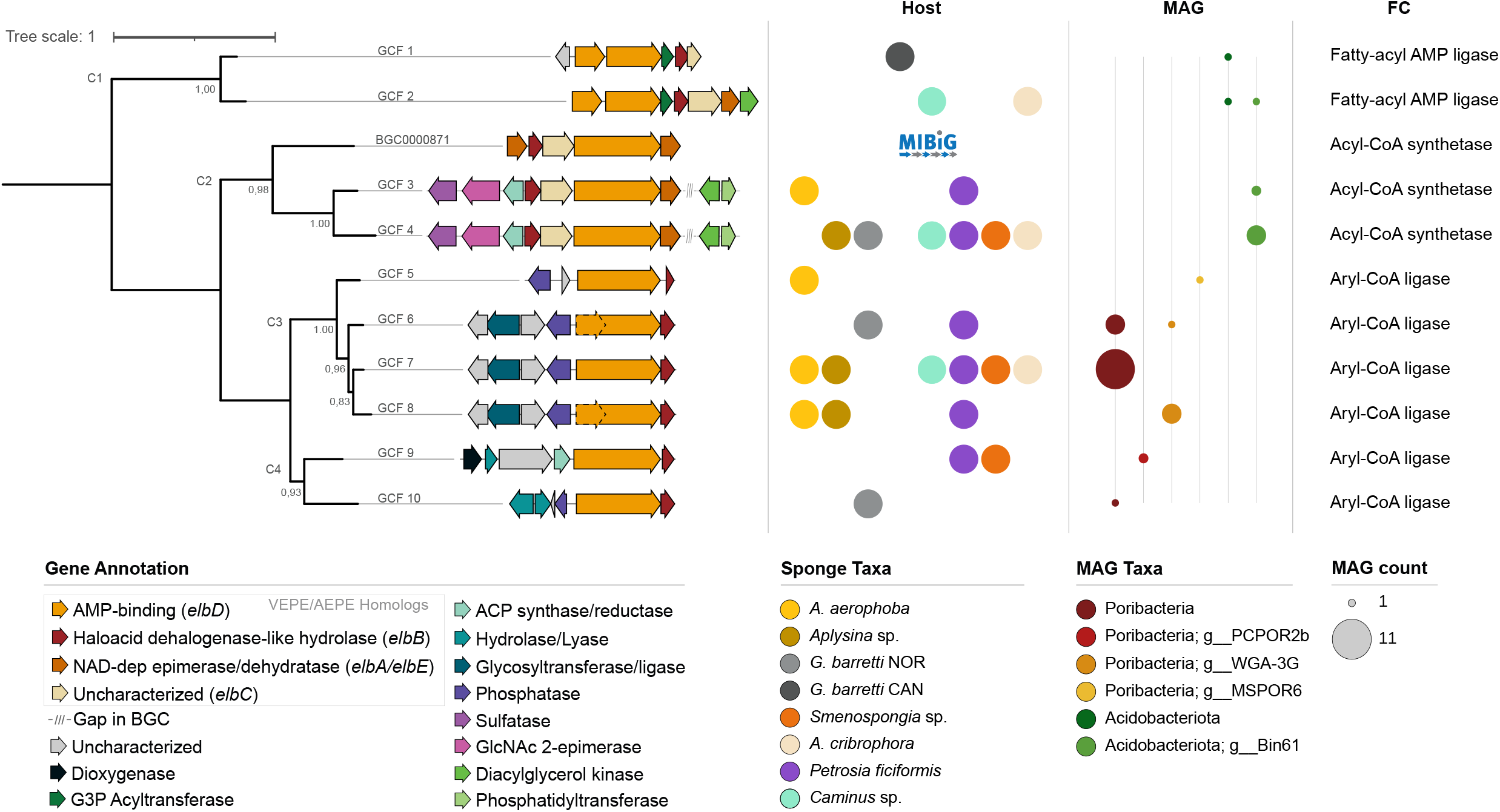
Diversity and characterization of GCFs putatively involved in EL production in sponge symbionts. FastTree phylogeny of GCFs putatively involved in EL production based on sequence similarity of the *elbD* homologue’s AMP-binding domain, rooted at midpoint, with local support values shown for internal nodes (range 0-1). Each GCF is depicted by its BiG-MAP representative. Genes are colored based on predicted function (PFAM domain), and dashed borders indicate a gene being present in some but not all members of a GCF. GCFs are annotated by their presence in host sponge species and in MAGs, where bubble size represents the number of distinct dereplicated MAGs for which the GCF is found. GCFs are also annotated by the AMP-binding domain’s predicted functional class (FC).

Searches for homologous genes coding for bacterial EL biosynthetic pathways were performed with NCBI tBLASTn using the characterized EL biosynthetic proteins; ElbD (UniProt ID: Q1DC43) and Agps (UniProt ID: Q1DBP5) from *Myxococcus xanthus*, the Ger (ENA accession: AGL49404) and the membrane-spanning lipid synthase (Mss) (ENA accession: AGL49750) from *Thermotoga maritima*, and the (putative) tetraether synthase (Tes) (NCBI accession: WP_015897912.1) from *Acidobaterium sp*. as queries. Hits were analyzed for domain architecture, as many had low percent amino acid identity. Hits, or contigs containing hits in the case of Ger, were annotated using Prodigal v2.6.3 (Hyatt *et al*., 2010) with standard parameters plus -p meta to obtain protein coding sequences. Domains were then annotated using HMMER3 v3.3 (Finn *et al*., 2011) using the hmmscan command with standard parameters and the Pfam-A database to scan for pfams present in the BLAST hits.

Searches for homologous genes coding for animal EL biosynthetic genes were performed with NCBI tBLASTn using the human AGPS (Uniprot ID: O00116.1) and plasmanylethanolamine desaturase (PEDS1) (UniProt ID: A5PLL7.3) proteins in the public genomes of *G. barretti* (Genbank accession: GCA_030341885.1), *A. aerophoba* (Genbank accession: GCA_949841015.1), and *P. ficiformis* (Genbank accession: GCA_947044365.1). *G. barretti* was searched on UniProt using the enzyme names “Plasmanylethanolamine desaturase” and “Alkylglycerone-phosphate synthase” to find the annotated proteins. Of the resulting *G. barretti* AGPS and PEDS1 hits, we considered only those which had the same InterPro domain composition as their human counterpart queries.

## Results and Discussion

### Lipid characterization in screened sponges confirms the presence of diverse ether lipids

Three sponge species: *Aplysina aerophoba, Geodia barretti and Petrosia ficiformis* were analyzed for their ether-lipid compositions. Alkyl ethers-based lipids are referred to here as Monoalkyl glycerol ethers (MGEs), while plasmalogens (vinyl, alkenyl ethers) are referred to here as dimethylacetals (DMAs), which are derived from the plasmalogens upon hydrolysis during the analysis process.

The main identified compounds were a range of MGEs with hydrocarbon chain lengths ranging from 14 to 16 (I.e., C_14:0_, C_15:0_, C_16:1_, C_16:0_) for all analyzed species. *A. aerophoba* also yielded longer chain compounds (i.e., the C17:0 and the C_18:0_ MGEs). Methylated MGEs (diMe-C_14:0_, Me-C_15:0_ and Me-C_16:0_) were detected in all samples, but the Me-C_18:0_ MGE was not found in *A. aerophoba*, and a Me-C_14:0_ was only present in *G. barretti*. Additionally, the *iso-*C_15:0_, *anteiso*-C_15:0_ and *iso-*C_16:0_ MGEs were detected in all the samples, and *A. aerophoba* also contained the *iso-*C_17:0_, *anteiso-*C_17:0_ and the *iso-*C_18:0_ MGE. We also identified membrane-spanning lipids MGE (i.e., diacids MGE) of C_30_ to C_31_ chain length in all the species, some of them containing saturated and methyl-branched variations (Supplementary Table 1).

DMA derived from plasmalogens were detected in all sponges, ranging from the C_22:0_ to C_26:0_ chain length, although not all were present in all the samples (Supplementary Table 1). The presence of plasmalogens with long carbon chains in bacteria is unusual as bacteria rarely synthesize fatty acids of more than 19 carbon atoms (Litchfield *et al*., 1976; de Kluijver *et al*., 2021). This suggests that the hydrocarbon chain of these plasmalogens is synthesized by the sponge host as previously reported for Demospongiae sponges (Dembitsky *et al*., 1989).

The lipid analysis confirmed the presence of both alkyl and plasmalogens (alkenyl) ether (as MGE and DMA, respectively) lipid compounds in all the studied sponges. Some small variations in the chain length and branching composition were observed between species. This diversity of chain length or methyl-branched compounds could be a result from specific adaptations of the sponge bacterial symbionts and/or host to its environment.

### Bacterial ether lipid gene clusters in marine sponge holobionts are diverse and widespread

#### Prediction of diverse elbB-elbE EL biosynthetic gene clusters

In order to investigate potential bacterial producers of the observed (and potentially unobserved) ether lipids, sponge metagenomes and MAGs were first queried using antiSMASH and compared to the MIBiG database. Previous characterization of the complete biosynthetic potential encoded in the marine sponge holobiont of three HMA species (Loureiro *et al*., 2022) had revealed several gene cluster families (GCFs) showing similarity to the MIBiG-BGC BGC0000871, the *elb* pathway described in *M. xanthus* (Lorenzen *et al*., 2014). Here, we expanded the knowledge on GCFs potentially involved in ether lipid production with an additional four HMA Demospongiae species. Manual selection and trimming of this initial set of BGCs yielded a final set of 92 BCGs, grouped into 10 GCFs, that excluded truncated BGCs and feature curated BGC borders. Based on GCF gene architectures, as well as A-domain phylogeny, we further grouped these BGCs into 4 clades (C1-C4, Fig. 2), of which clade 1 had not been previously reported.

We observed a variable distribution of GCFs across host sponge species (Fig. 2), with both widespread GCFs, i.e. GCFs 4 and 7 being present in six of the seven host sponge species, and sponge specific GCFs, i.e. GCFs 1, 5 and 10 present in a single sponge species. The majority of the sponge holobionts (*G. barretti* NOR, *Aciculites cribrophora, Smenospongia* sp., *Caminus* sp. and *P. ficiformis*) show high potential diversity in ether lipid biosynthetic profiles, harboring GCFs belonging to three of the four clades. *G. barretti* CAN shows the least diverse profile, featuring a single GCF/clade. *G. barretti* NOR (Norway) samples were collected up to approximately 500m depth, in contrast to *G. barretti* CAN (Canada) samples collected below 1000m depth (Table S1). A recent study showed that several *G. barretti*-derived bioactive compounds, as well as *G. barretti* bacterial symbionts, were detected exclusively above 1000 m depth (Steffen *et al*., 2022). This study uncovers an analogous dichotomy, with the mutually exclusive presence of both GCFs and MAGs recovered from *G. barretti* CAN & NOR.

Seventy two (78%) of the *elb*-like BGCs were recovered in MAGs belonging to the candidate phylum Poribacteria and the phylum Acidobacteriota (Subdivision 6). The remaining 22% were not present in MAGs. Additionally, querying NCBI’s nr database using Cblaster (Gilchrist *et al*., 2021) with BGC0000871 (the *elbB-elbE* BGC) as well as a GCF 4 member (given its similarity to BGC0000871) yielded hits belonging to over 150 organisms, the majority of which belong to the Acidobacteriota and Proteobacteria phyla (Fig. S1,S2). This further supports the potential synthesis of ELs by this bacterial phylum in sponges, and points towards a more ubiquitous presence in other bacterial groups than what has been previously reported for these BGCs. Furthermore, GCFs shared across a higher number of sponge hosts also appear to be present in more distinct dereplicated MAGs, as is the case for GCF 7 (Fig. 2). This could be an indication of parallel evolution and diversification of these MAGs in each of the host sponge species.

The architectural features of each GCF (Fig. 2), as well as host MAG phylogeny (Fig. S5), are congruent with the phylogenetic clades generated based on *elbD*’s A-domain sequence similarity, an indication of BGC evolutionary conservation. An exception to this congruence is seen for GCFs 6 and 8, which include BGCs with a canonical *elbD*, as well as a variant in which the *elbD* homolog is split into two open reading frames. In these cases, the split is made between the Re and A domains. Contrarily to their placement in separate GCFs by BiG-SCAPE, a BGC clustering tool which considers BGC gene and protein domain architecture (Navarro-Muñoz *et al*., 2020), BGCs with a split *elbD* gene cluster together based on both A-domain sequence similarity (Fig. S4) and host MAG phylogeny (Fig. S3). While this split variant is present only in MAGs of poribacterial origin in our dataset, soil-derived acidobacterial homologs of this variant have also been reported (Sinninghe Damsté *et al*., 2018). The BGCs from these GCFs are present in two distinct poribacterial taxa. In several cases, both variants and MAGs coexist in the same host sponge sample (Fig. S3), which is in line with previous findings that different Poribacteria taxonomic clades can coexist within the same sponge host (86). This may suggest divergent evolution of the variants, and/or complementation of loss of function of one of the variants.

The core gene architectures (i.e., the set of genes homologous to key enzyme-coding genes from the *elb*-BGC) of these putatively EL-associated GCFs also display a gradient in diversity, with GCFs 3 and 4 harboring four of the five core genes in the same orientation as the original *elb*-BGC,, and GCFs 5-10 harboring only *elbD* and *elbB* homologs (Figure 2). Analysis of the A-domain active site specificity-conferring residues (using AdenylPred v1 (Robinson *et al*., 2020) and PARAS v0.0.3 ((Terlouw et al. 2025), Table S4) indicates potentially different functional classes and substrate specificities for Clade 1, Clade 2 and Clades 3-4. It must be noted that the prediction scores of both tools on these queries are poor, likely because no sponge-holobiont-derived A-domain sequences are present in the model training datasets. Nevertheless, together with these predictions, differences in the active site residues that are known to determine substrate specificities suggest additional chemical diversity in the molecules produced by these GCFs (Table S4). Additionally, the co-occurrence of distinct MAGs containing BGCs from several GCFs in the same sponge metagenomic sample also supports the hypothesis that these BGCs are responsible for the production of chemically and functionally diverse molecules.

Domain annotations of genes adjacent to *elbD* and/or *elbB* are also variable across the dataset (Fig. 2). Clades 1 and 2 feature adjacent genes that are tightly connected to phospholipid biosynthesis which could play a role in ether lipid precursor biosynthesis, as glycerol-3-phosphate acyltransferases and diacylglycerol (DAG) kinases are involved in the synthesis of phosphatidic acid, the main membrane phospholipid precursor (van Blitterswijk and Houssa, 2000; Li *et al*., 2017), and cytidine-5⍰-diphosphate (CDP)-alcohol phosphatidyltransferases are involved in the synthesis of polar head groups from CDP-DAG (Centola *et al*., 2021). Clades 2 and 4 additionally feature 3-oxoacyl-[acyl-carrier-protein (ACP)] synthases and enoyl-ACP reductases which are both involved in fatty acid biosynthesis (Rock and Cronan, 1996). Clade 2 also features arylsulfatases and an N-acylglucosamine 2-epimerase (GlcNAc 2-epimerase). These bidirectional enzymes could participate in the production of sulfated ELs similar to the sulfated ether glycoglycerolipids that have been shown to be produced by coral symbionts (Dodgson *et al*., 1956; Ghosh and Roseman, 1965; Sikorskaya *et al*., 2021). Clade 3 features a phosphatase which could be involved in additional modification of phospholipids, as well as lipopolysaccharide-related glycosyltransferases that could be part of the synthesis of ether-linked glycoglycerolipids. Such ether glycoglycerolipids have been previously reported in marine sponges (Genin *et al*., 2004), and Poribacteria have been predicted, based on genomics, to incorporate phosphoinositol-linked glycolipids into their cell membranes (Kamke *et al*., 2013; Podell *et al*., 2019). Finally, several of the GCFs feature genes coding for transporters and efflux systems further up/downstream of the BGC core, which could be involved in export of molecules being used in signaling or defense mechanisms (Table S3).

Nevertheless, future studies are needed to assess the ecophysiological advantages and conditions leading to the biosynthetic diversity of EL-BGCs in these bacterial phyla, and their role within the sponge holomicrobiome.

#### Targeted genomic prediction of alternative EL biosynthetic pathways in bacteria

The biosynthetic genomic investigations performed above focused on clustered specialized metabolism BGCs, thus lacking the ability to target standalone proteins that may play a role in EL production. We therefore performed targeted genomic searches using the key genes of the aerobic and anaerobic EL pathways for alkyl and plasmalogen (alkenyl) ethers (i.e., *elbD, agps, ger*, see introduction for details). Additionally, we included in these queries the membrane-spanning lipid (MSL) synthase gene *mss*, involved in the coupling of fatty acids leading to MSLs (with or without ether bonds), as well as the *tes* gene homologs of which have been proposed to catalyze the reaction of coupling of fatty acids in bacterial MSLs, that have not been experimentally validated (Sahonero Canavesi et al., 2022; Zeng et al., 2022).

Sequence homologs to the *elbD* and *agps* genes of *M. xanthus* were detected in all sponge metagenomes analyzed. *elbD* hits were detected in Poribacteria and Acidobacteriota MAGs, as both standalone *elbD* hits and as part of *elb*-BGCs (supplementary table 2). Standalone *elbD* hits were additionally found in Latescibacterota MAGs, a taxonomic group that, to our knowledge, has not yet been associated with the capacity to produce ELs. Some *agps* gene hits were co-localized in the same MAGs containing *elbD* hits, as observed in the case of *M. xanthus* (Lorenzen et al., 2014) (supplementary table 2). The co-localized *elbD* and *agps* MAGs belonged to Poribacteria, Acidobacteriota, and Latescibacterota. These results suggest that at least some members of these three phyla, present in all species of the analyzed sponges, have the genomic capacity to synthesize ELs, both in the form of alkyl ethers (via the *elbD* gene/pathway), and/or in the form of plasmalogens (via the *agps*). However, no short-chain plasmalogens (generally associated with bacterial origin) were detected in our lipid analysis, so it is possible that bacterial *agps* were not expressed at the time of collection.

When searching for the possibility of anaerobic EL production via Ger homologs (Jackson et al., 2020) (Fig. 1), no full-length hits were found using a query of *ger* from *Thermotoga maritima* in any metagenome or MAG.

However, as *ger* comprises both *plsA* and *plsR*, some hits to the *plsA* portion of the query were obtained (Supplementary table 3). The majority of the hits contained an ATPase/Co-A activase domain (PF01869) in duplicate and a reduction domain (PF09989), as seen in *ger* architecture 2 (Sahonero-Canavesi *et al*., 2022) (Supplementary Table 3). The *ger* homologs were found in MAGs belonging to members of the Chloroflexota, Acidobacteriota, Proteobacteria, and Latescibacterota phyla. Domain analysis of the *ger* homologs suggest these might be involved in the formation of alkyl ethers (by Ger) since the second reduction domain associated with the plsA/plsR plasmalogen biosynthesis is not present in those hits (Sahonero-Canavesi *et al*., 2022), but these proteins have been seen to be only active under anaerobic conditions (Sahonero-Canavesi *et al*., 2022).

Thus, these results support the possible synthesis of ELs in sponges under different oxygen concentrations. Although sponges are known to take up oxygen by pumping in oxygenated seawater, they are quite resilient to hypoxia (Micaroni *et al*., 2022; Riisgård, 2024) and both *A. aerophoba* and *G. barretti* have been shown to harbor anoxic micro-environments (Hoffmann *et al*., 2005, 2008). It is possible that anoxic niches formed within the tissue are compatible with facultative anaerobic metabolism of members of the Acidobacteriota, Chloroflexota, Proteobacteria and Latescibacterota, and of the synthesis of ELs under anaerobic conditions. This is the first time that the formation of ELs in sponges is linked to these taxonomic groups. Formation of alkyl ethers by the Ger homolog under anaerobic conditions cannot be discarded, as alkyl ethers in general have been detected in the lipid analysis but it is not possible to discern here if these were synthesized by the aerobic/facultative *elb*-BGC or by the anaerobic Ger pathway. We found that all sponge species had at least one MAG with *mss/tes* and *elbD/ger* (supplementary table 4). All MAGs with *mss* co-occurred only with *elbD* and belonged to the Poribacteria. Accordingly, all MAGs with *tes* co-occurred with *elbD* or *ger* and belonged to the Acidobacteriota, suggesting that both bacteria phyla have the potential to produce the membrane-spanning MGEs using distinct pathways.

The *elb*-pathway analysis along with the targeted alternative pathway analysis show that sponge symbionts harbor the potential to synthesize ELs by using different pathways under different oxygen conditions. This adaptive strategy is especially useful for sponge symbionts as the micro-environments in the host tissues can be subjected to multiple environmental pressures and oxygen concentrations. Likewise, sponges themselves may select for such EL producers and thus, having abundant ELs could grant the host sponge with niche colonizing advantages.

### Biosynthetic capacity of the sponge host to produce ether lipids

In the performed lipid analysis, three out of the four types of detected ether lipids (Supplementary table 1) can be attributed to bacterial members of the sponge holobiont, given their chain length, the presence of mid-chain fatty acids (i.e., normally attributed to bacteria; REG), and the ether lipid biosynthesis genes found in the performed genomic analysis. However, the detected plasmalogens (DMAs) with C_20_-C_27_ carbon chains are more likely attributed to the sponge host on the basis of their longer carbon chain length (REFs).

To investigate whether the sponge host has the capacity to synthesize these long chain plasmalogens, we searched for the eukaryotic genes encoding for the proteins responsible for plasmalogen ether lipid biosynthesis, *agps* and plasmanylethanolamine desaturase (*PEDS1*), in the published genomes of *G. barretti, A. aerophoba*, and *P. ficiformis*.

At the time of the analysis, only the recently released *G. barretti* genome (Steffen *et al*., 2023) had full protein annotations in the UniProt database. When searching UniProt for AGPS and PEDS1 protein homologues in *G. barretti*, we found one protein entry annotated as AGPS (UniProtKB ID: A0AA35RJ59), as well as one protein annotated as PEDS1 (UniProtKB ID: A0AA35WZZ7), both featuring domain architecture consistent with that of its human counterparts, lending confidence to these hits.

For the unannotated *A. aerophoba* and *P. ficiformis* genomes, we performed an homology-based search using tblastn with human AGPS and PEDS1 protein sequences as queries. We detected hits in both genomes with high amino acid sequence homology to human AGPS and PEDS1 proteins (supplementary table 5). Given the genomic evidence that these proteins exist in all three sponge species, it is likely that the host itself can produce the long-chain ether lipid plasmalogens we detected in these species.

## Conclusion

Alkyl and alkenyl (plasmanogen) ether lipids have been described in the marine sponge holobiont, but so far, their biosynthetic pathways and producers remained elusive. Here, we used biosynthetic genomic searches in sponge symbiont MAGs and public sponge genomes to elucidate the biosynthetic origins of observed ELs in diverse HMA sponges. A diverse set of GCFs with similarity to the validated *elb*-BGC was present in MAGs belonging to Acidobacteriota and Poribacteria that putatively code for the proteins involved in producing the detected MGEs, methyl-MGEs, and membrane-spanning MGEs. Latescibacterota and Chloroflexota also emerged as potential EL producers, specifically of the observed MGEs, given their possession of ElbD and/or Ger homologs, and these phyla have not previously been connected with the ability to make ELs. Sponge genomes harbor genes related to the observed long chain DMAs, thus accounting for all the types of ELs detected in the sponges investigated here. Illuminating which sponge symbionts have the ability to produce ELs and the pathways they use to do so can aid in the search for biotechnologically relevant molecules, start to define the elusive roles of symbionts in the sponge holobionts, and possibly lead to the discovery of novel enzymes or pathways for EL production.

## Acknowledgments

The authors wish to thank Ellen Kenchington for the Canadian *G. barretti* samples, the late Hans Tore Rapp for his repeated help to sample the Norwegian fjord *G. barretti* specimens, and Adriaan Schrier for offering his submarine to collect sponges in Dominica. We thank Caglar Yildiz for valuable advice and discussions. We thank Michel Koenen for the processing and interpretation of lipid analyses.

This research was financially supported by the VLAG NWO PhD project “*Mare incognita*” to CL; the European Commission through Horizon2020 project SponGES (Grant agreement ID: 679849) to DS, and AG; the Marie Sklodowska-Curie Individual Fellowship COSMos (Grant agreement ID: 897121) and a Wageningen Graduate Schools Postdoc Talent Programme grant to MAS; and the SIAM Gravitation Grant (024.002.002) from the Dutch Ministry of Education, Culture and Science (OCW) to LV and DXSC.

## Author contributions

C.L: Conceptualization, data curation, formal analysis, funding acquisition, investigation, methodology, software, visualization, writing - original draft, writing - review & editing.

M.A.S: Conceptualization, data curation, formal analysis, funding acquisition, investigation, methodology, software, writing - original draft, writing - review & editing.

D.X.S.C.: Data curation, formal analysis, investigation, methodology, visualization, writing - review & editing.

A.G: Data curation, investigation, writing - review & editing.

V.G.: Resources.

J.vdO: Supervision, funding acquisition, writing - review & editing.

L.V.: Supervision, funding acquisition, writing - review & editing.

M.H.M: Conceptualization, funding acquisition, supervision, writing - review & editing.

D.S: Conceptualization, funding acquisition, resources, supervision, writing - review & editing.

## Data availability

The data for this study have been deposited in the European Nucleotide Archive (ENA) at EMBL-EBI under accession numbers PRJEB51534, PRJEB59408. Python scripts created for this analysis are available at https://github.com/CatarinaCarolina/sponge_ether_lipid. Figure S5: the complete BGC A-domain based phylogeny iTOL tree, complementary to Fig. S4, can be found here: https://itol.embl.de/tree/62195245151491291644849392.

## Ethics Statement

## Figures

**Figure 1:** (needs to be simplified still) fig1_combined_ac.pdf

**Figure 2:** fig2_itol_vepe_domexp_c03_tg01_c50_nonames_addbins_bs.pdf

Appendix Tables: Supplementary tables v5.xlsx

## References

Agarwal, V., Blanton, J.M., Podell, S., Taton, A., Schorn, M.A., Busch, J., et al. (2017) Metagenomic discovery of polybrominated diphenyl ether biosynthesis by marine sponges. Nat Chem Biol 13: 537–543.

Alam, N., Bae, B.H., Hong, J., Lee, C.O., Shin, B.A., Im, K.S., and Jung, J.H. (2001) Additional bioactive Lyso-PAF congeners from the sponge Spirastrella abata. J Nat Prod 64: 533–535.

Ben-Califa, N., Bishara, A., Kashman, Y., and Neumann, D. (2012) Salarin C, a member of the salarin superfamily of marine compounds, is a potent inducer of apoptosis. Invest New Drugs 30: 98–104.

Blin, K., Shaw, S., Steinke, K., Villebro, R., Ziemert, N., Lee, S.Y., et al. (2019) antiSMASH 5.0: updates to the secondary metabolite genome mining pipeline. Nucleic Acids Res 47: W81–W87.

van Blitterswijk, W.J. and Houssa, B. (2000) Properties and functions of diacylglycerol kinases. Cell Signal 12: 595–605.

Braverman, N.E. and Moser, A.B. (2012) Functions of plasmalogen lipids in health and disease. Biochim Biophys Acta 1822: 1442–1452.

Calcabrini, C., Catanzaro, E., Bishayee, A., Turrini, E., and Fimognari, C. (2017) Marine Sponge Natural Products with Anticancer Potential: An Updated Review. Mar Drugs 15.:

Carroll, A.R., Copp, B.R., Davis, R.A., Keyzers, R.A., and Prinsep, M.R. (2019) Marine natural products. Nat Prod Rep 36: 122–173.

Centola, M., van Pee, K., Betz, H., and Yildiz, Ö. (2021) Crystal structures of phosphatidyl serine synthase PSS reveal the catalytic mechanism of CDP-DAG alcohol O-phosphatidyl transferases. Nat Commun 12: 6982.

Chaib De Mares, M., Jiménez, D.J., Palladino, G., Gutleben, J., Lebrun, L.A., Muller, E.E.L., et al. (2018) Expressed protein profile of a Tectomicrobium and other microbial symbionts in the marine sponge Aplysina aerophoba as evidenced by metaproteomics. Sci Rep 8: 11795.

Dembitsky, V.M., Gorina, I.A., Fedorova, I.P., and Solovieva, M.V. (1989) Comparative investigation of plasmalogens, alkylacyl and diacyl glycerophospholipids of the marine sponges (type Porifera, class Demospongiae). Comparative Biochemistry and Physiology Part B: Comparative Biochemistry 92: 733–736.

Djerassi, C. and Lam, W.K. (1991) Phospholipid studies of marine organisms. Part 25. Sponge phospholipids. Acc Chem Res 24: 69–75.

Dodgson, K.S., Spencer, B., and Williams, K. (1956) Studies on sulphatases. 13. The hydrolysis of substituted phenyl sulphates by the arylsulphatase of Alcaligenes metalcaligenes. Biochem J 64: 216–221.

Dyshlovoy, S.A., Fedorov, S.N., Svetashev, V.I., Makarieva, T.N., Kalinovsky, A.I., Moiseenko, O.P., et al. (2022) 1-O-Alkylglycerol Ethers from the Marine Sponge Guitarra abbotti and Their Cytotoxic Activity. Mar Drugs 20.:

Edgar, R.C. (2004) MUSCLE: multiple sequence alignment with high accuracy and high throughput. Nucleic Acids Res 32: 1792–1797.

Engelberts, J.P., Robbins, S.J., de Goeij, J.M., Aranda, M., Bell, S.C., and Webster, N.S. (2020) Characterization of a sponge microbiome using an integrative genome-centric approach. ISME J 14: 1100–1110.

Essack, M., Bajic, V.B., and Archer, J.A.C. (2011) Recently confirmed apoptosis-inducing lead compounds isolated from marine sponge of potential relevance in cancer treatment. Mar Drugs 9: 1580–1606.

Fan, L., Reynolds, D., Liu, M., Stark, M., Kjelleberg, S., Webster, N.S., and Thomas, T. (2012) Functional equivalence and evolutionary convergence in complex communities of microbial sponge symbionts. Proc Natl Acad Sci U S A 109: E1878–87.

Fedorov, S.N., Makarieva, T.N., Guzii, A.G., Shubina, L.K., Kwak, J.Y., and Stonik, V.A. (2009) Marine two-headed sphingolipid-like compound rhizochalin inhibits EGF-induced transformation of JB6 P+ Cl41 cells. Lipids 44: 777–785.

Finn, R.D., Clements, J., and Eddy, S.R. (2011) HMMER web server: interactive sequence similarity searching. Nucleic Acids Res 39: W29–37.

Gallego-García, A., Monera-Girona, A.J., Pajares-Martínez, E., Bastida-Martínez, E., Pérez-Castaño, R., Iniesta, A.A., et al. (2019) A bacterial light response reveals an orphan desaturase for human plasmalogen synthesis. Science 366: 128–132.

Genin, E., Njinkoue, J.M., Wielgosz-Collin, G., Houssay, C., and Barnathan, G. (2004) Glycolipids from marine sponges: Monoglycosylceramides and alkyldiglycosylglycerols: Isolation, characterization and biological activity. 68: 327–334.

Ghosh, S. and Roseman, S. (1965) THE SIALIC ACIDS. V. N-ACYL-D-GLUCOSAMINE 2-EPIMERASE. J Biol Chem 240: 1531–1536.

Gilchrist, C.L.M., Booth, T.J., van Wersch, B., van Grieken, L., Medema, M.H., and Chooi, Y.-H. (2021) cblaster: a remote search tool for rapid identification and visualization of homologous gene clusters. Bioinform Adv 1: vbab016.

Goldfine, H. (2022) Plasmalogens in bacteria, sixty years on. Front Mol Biosci 9: 962757.

Goldfine, H. (2017) The anaerobic biosynthesis of plasmalogens. FEBS Lett 591: 2714–2719.

Gramzow, M., Schröder, H.C., Fritsche, U., Kurelec, B., Robitzki, A., Zimmermann, H., et al.(1989) Role of phospholipase A2 in the stimulation of sponge cell proliferation by homologous lectin. Cell 59: 939–948.

Hentschel, U., Fieseler, L., Wehrl, M., Gernert, C., Steinert, M., Hacker, J., and Horn, M. (2003) Microbial Diversity of Marine Sponges. In Sponges (Porifera). Müller, W.E.G. (ed). Berlin, Heidelberg: Springer Berlin Heidelberg, pp. 59–88.

Hoffmann, F., Larsen, O., Thiel, V., Rapp, H.T., Pape, T., Michaelis, W., and Reitner, J. (2005) An Anaerobic World in Sponges. Geomicrobiol J 22: 1–10.

Hoffmann, F., Røy, H., Bayer, K., Hentschel, U., Pfannkuchen, M., Brümmer, F., and de Beer, D. (2008) Oxygen dynamics and transport in the Mediterranean sponge Aplysina aerophoba. Mar Biol 153: 1257–1264.

Hyatt, D., Chen, G.-L., Locascio, P.F., Land, M.L., Larimer, F.W., and Hauser, L.J. (2010) Prodigal: prokaryotic gene recognition and translation initiation site identification. BMC Bioinformatics 11: 119.

Jackson, D.R., Cassilly, C.D., Plichta, D.R., Vlamakis, H., Liu, H., Melville, S.B., et al. (2021) Plasmalogen Biosynthesis by Anaerobic Bacteria: Identification of a Two-Gene Operon Responsible for Plasmalogen Production in Clostridium perfringens. ACS Chem Biol 16: 6–13.

Kamke, J., Rinke, C., Schwientek, P., Mavromatis, K., Ivanova, N., Sczyrba, A., et al. (2014) The candidate phylum Poribacteria by single-cell genomics: new insights into phylogeny, cell-compartmentation, eukaryote-like repeat proteins, and other genomic features. PLoS One 9: e87353.

Kamke, J., Sczyrba, A., Ivanova, N., Schwientek, P., Rinke, C., Mavromatis, K., et al. (2013) Single-cell genomics reveals complex carbohydrate degradation patterns in poribacterial symbionts of marine sponges. ISME J 7: 2287–2300.

Kielak, A.M., Barreto, C.C., Kowalchuk, G.A., van Veen, J.A., and Kuramae, E.E. (2016) The Ecology of Acidobacteria: Moving beyond Genes and Genomes. Front Microbiol 7: 744.

de Kluijver, A., Nierop, K.G.J., Morganti, T.M., Bart, M.C., Slaby, B.M., Hanz, U., et al. (2021) Bacterial precursors and unsaturated long-chain fatty acids are biomarkers of North-Atlantic deep-sea demosponges. PLoS One 16: e0241095.

Letunic, I. and Bork, P. (2019) Interactive Tree Of Life (iTOL) v4: recent updates and new developments. Nucleic Acids Res 47: W256–W259.

Litchfield, C., Greenberg, A.J., Noto, G., and Morales, R.W. (1976) Unusually high levels of C24-C30 fatty acids in sponges of the class Demospongiae. Lipids 11: 567–573.

Li, Z., Tang, Y., Wu, Y., Zhao, S., Bao, J., Luo, Y., and Li, D. (2017) Structural insights into the committed step of bacterial phospholipid biosynthesis. Nat Commun 8: 1691.

Lorenzen, W., Ahrendt, T., Bozhüyük, K.A.J., and Bode, H.B. (2014) A multifunctional enzyme is involved in bacterial ether lipid biosynthesis. Nat Chem Biol 10: 425–427.

Loureiro, C., Galani, A., Gavriilidou, A., de Mares, M.C., van der Oost, J., Medema, M.H., and Sipkema, D. (2022) Comparative metagenomic analysis of biosynthetic diversity across sponge microbiomes highlights metabolic novelty, conservation and diversification. bioRxiv 2022.04.01.486688.

McMurray, S.E., Johnson, Z.I., Hunt, D.E., Pawlik, J.R., and Finelli, C.M. (2016) Selective feeding by the giant barrel sponge enhances foraging efficiency. Limnol Oceanogr 61: 1271–1286.

Medema, M.H., Kottmann, R., Yilmaz, P., Cummings, M., Biggins, J.B., Blin, K., et al. (2015) Minimum Information about a Biosynthetic Gene cluster. Nat Chem Biol 11: 625–631.

Micaroni, V., Strano, F., McAllen, R., Woods, L., Turner, J., Harman, L., and Bell, J.J. (2022) Adaptive strategies of sponges to deoxygenated oceans. Glob Chang Biol 28: 1972–1989.

Mohsenian Kouchaksaraee, R., Moridi Farimani, M., Li, F., Nazemi, M., and Tasdemir, D. (2020) Integrating Molecular Networking and 1H NMR Spectroscopy for Isolation of Bioactive Metabolites from the Persian Gulf Sponge Axinella sinoxea. Mar Drugs 18.:

Morita, M. and Schmidt, E.W. (2018) Parallel lives of symbionts and hosts: chemical mutualism in marine animals. Nat Prod Rep 35: 357–378.

Müller, W.E.G., Klemt, M., Thakur, N.L., Schröder, H.C., Aiello, A., D’Esposito, M., et al. (2004) Molecular/chemical ecology in sponges: evidence for an adaptive antibacterial response in Suberites domuncula. Mar Biol 144: 19–29.

Mu, R., Momeni, S., Krieger, M., Xie, B., Cao, X., Merritt, J., and Wu, H. (2024) Plasmalogen, a glycerophospholipid crucial for Streptococcus mutans acid tolerance and colonization. Appl Environ Microbiol e0150023.

Navarro-Muñoz, J.C., Selem-Mojica, N., Mullowney, M.W., Kautsar, S.A., Tryon, J.H., Parkinson, E.I., et al. (2020) A computational framework to explore large-scale biosynthetic diversity. Nat Chem Biol 16: 60–68.

Olm, M.R., Brown, C.T., Brooks, B., and Banfield, J.F. (2017) dRep: a tool for fast and accurate genomic comparisons that enables improved genome recovery from metagenomesthrough de-replication. ISME J 11: 2864–2868.

Olm, M.R., Crits-Christoph, A., Diamond, S., Lavy, A., Matheus Carnevali, P.B., and Banfield, J.F. (2020) Consistent Metagenome-Derived Metrics Verify and Delineate Bacterial Species Boundaries. mSystems 5.:

Pascal Andreu, V., Augustijn, H.E., van den Berg, K., van der Hooft, J.J.J., Fischbach, M.A., and Medema, M.H. (2021) BiG-MAP: an Automated Pipeline To Profile Metabolic Gene Cluster Abundance and Expression in Microbiomes. mSystems 6: e0093721.

Peters, E.E., Cahn, J.K.B., Lotti, A., Gavriilidou, A., Steffens, U.A.E., Loureiro, C., et al. (2023) Distribution and diversity of “Tectomicrobia”, a deep-branching uncultivated bacterial lineage harboring rich producers of bioactive metabolites. ISME Commun 3: 50.

Podell, S., Blanton, J.M., Neu, A., Agarwal, V., Biggs, J.S., Moore, B.S., and Allen, E.E. (2019) Pangenomic comparison of globally distributed Poribacteria associated with sponge hosts and marine particles. ISME J 13: 468–481.

Price, M.N., Dehal, P.S., and Arkin, A.P. (2010) FastTree 2--approximately maximum-likelihood trees for large alignments. PLoS One 5: e9490.

Quévrain, E., Barnathan, G., Meziane, T., Domart-Coulon, I., Rabesaotra, V., and Bourguet-Kondracki, M.-L. (2012) New 2-methyl-13-icosenoic acid from the temperate calcisponge Leuconia johnstoni. Lipids 47: 345–353.

Quinn, R.A., Vermeij, M.J.A., Hartmann, A.C., Galtier d’Auriac, I., Benler, S., Haas, A., et al. (2016) Metabolomics of reef benthic interactions reveals a bioactive lipid involved in coral defence. Proc Biol Sci 283.:

Řezanka, T., Křesinová, Z., Kolouchová, I., and Sigler, K. (2012) Lipidomic analysis of bacterial plasmalogens. Folia Microbiol 57: 463–472.

Riisgård, H.U. (2024) Oxygen Extraction Efficiency and Tolerance to Hypoxia in Sponges. J Mar Sci Eng 12: 138.

Rix, L., Ribes, M., Coma, R., Jahn, M.T., de Goeij, J.M., van Oevelen, D., et al. (2020) Heterotrophy in the earliest gut: a single-cell view of heterotrophic carbon and nitrogen assimilation in sponge-microbe symbioses. ISME J 14: 2554–2567.

Robinson, S.L., Terlouw, B.R., Smith, M.D., Pidot, S.J., Stinear, T.P., Medema, M.H., and Wackett, L.P. (2020) Global analysis of adenylate-forming enzymes reveals β-lactone biosynthesis pathway in pathogenic Nocardia. J Biol Chem 295: 14826–14839.

Rock, C.O. and Cronan, J.E. (1996) Escherichia coli as a model for the regulation of dissociable (type II) fatty acid biosynthesis. Biochim Biophys Acta 1302: 1–16.

Rubio-Portillo, E., Martin-Cuadrado, A.B., Caraballo-Rodríguez, A.M., Rohwer, F., Dorrestein, P.C., and Antón, J. (2020) Virulence as a Side Effect of Interspecies Interaction in Vibrio Coral Pathogens. MBio 11.:

Sahonero-Canavesi, D.X., Siliakus, M.F., Abdala Asbun, A., Koenen, M., von Meijenfeldt, F.A.B., Boeren, S., et al. (2022) Disentangling the lipid divide: Identification of key enzymes for the biosynthesis of membrane-spanning and ether lipids in Bacteria. Sci Adv 8: eabq8652.

Schorn, M.A., Jordan, P.A., Podell, S., Blanton, J.M., Agarwal, V., Biggs, J.S., et al. (2019) Comparative Genomics of Cyanobacterial Symbionts Reveals Distinct, Specialized Metabolism in Tropical Dysideidae Sponges. MBio 10.:

Sikorskaya, T.V., Efimova, K.V., and Imbs, A.B. (2021) Lipidomes of phylogenetically different symbiotic dinoflagellates of corals. Phytochemistry 181: 112579.

Sinninghe Damsté, J.S., Rijpstra, W.I.C., Foesel, B.U., Huber, K.J., Overmann, J., Nakagawa, S., et al. (2018) An overview of the occurrence of ether-and ester-linked iso-diabolic acid membrane lipids in microbial cultures of the Acidobacteria: Implications for brGDGT paleoproxies for temperature and pH. Org Geochem 124: 63–76.

Sinninghe Damsté, J.S., Rijpstra, W.I.C., Hopmans, E.C., Foesel, B.U., Wüst, P.K., Overmann, J., et al. (2014) Ether-and ester-bound iso-diabolic acid and other lipids in members of acidobacteria subdivision 4. Appl Environ Microbiol 80: 5207–5218.

Steffen, K., Indraningrat, A.A.G., Erngren, I., Haglöf, J., Becking, L.E., Smidt, H., et al. (2022) Oceanographic setting influences the prokaryotic community and metabolome in deep-sea sponges. Sci Rep 12: 3356.

Steffen, K., Proux-Wéra, E., Soler, L., Churcher, A., Sundh, J., and Cárdenas, P. (2023) Whole genome sequence of the deep-sea sponge Geodia barretti (Metazoa, Porifera, Demospongiae). G3 13.:

Thomas, T., Moitinho-Silva, L., Lurgi, M., Björk, J.R., Easson, C., Astudillo-García, C., et al. (2016) Diversity, structure and convergent evolution of the global sponge microbiome. Nat Commun 7: 1–12.

Uritskiy, G.V., DiRuggiero, J., and Taylor, J. (2018) MetaWRAP-a flexible pipeline for genome-resolved metagenomic data analysis. Microbiome 6: 158.

Van Soest, R.W.M., Boury-Esnault, N., Vacelet, J., Dohrmann, M., Erpenbeck, D., De Voogd, N.J., et al. (2012) Global diversity of sponges (Porifera). PLoS One 7: e35105.

Vítová, M., Palyzová, A., and Řezanka, T. (2021) Plasmalogens - Ubiquitous molecules occurring widely, from anaerobic bacteria to humans. Prog Lipid Res 83: 101111.

Webster, N.S. and Thomas, T. (2016) The Sponge Hologenome. MBio 7: e00135–16.

Zeng, Z., Chen, H., Yang, H., Chen, Y., Yang, W., Feng, X., Pei, H., Welander, P.V. (2022) Identification of a protein responsible for the synthesis of archaeal membrane-spanning GDGT lipids. Nat Commun 13: 1545.

